# A new massively parallel nanoball sequencing platform for whole exome research

**DOI:** 10.1101/451641

**Authors:** Yu Xu, Zhe Lin, Chong Tang, Yujing Tang, Yue Cai, Hongbin Zhong, Xuebin Wang, Wenwei Zhang, Chongjun Xu, Jingjing Wang, Jian Wang, Huanming Yang, Linfeng Yang, Qiang Gao

## Abstract

**Background:** Whole exome sequencing (WES) has been widely used in human genetics research. BGISEQ-500 is a recently established next-generation sequencing platform. However, the performance of BGISEQ-500 on WES is not well studied. In this study, we evaluated the performance of BGISEQ-500 on WES by side-to-side comparison with Hiseq4000, on well-characterized human sample NA12878.

**Results:** BGISEQ demonstrated similarly high reproducibility as Hiseq for variation detection. Also, the SNPs from BGISEQ data is highly consistent with Hiseq results (concordance 96.5%∼97%). Variation detection accuracy was subsequently evaluated with data from the genome in a bottle project as the benchmark. Both platforms showed similar sensitivity and precision in SNP detection. While in indel detection, BGISEQ showed slightly higher sensitivity and lower precision. The impact of sequence depth and read length on variation detection accuracy was further analyzed, and showed that variation detection sensitivity still increasing when the sequence depth is larger than 100x, and the impact of read length is minor when using 100x data.

**Conclusions:** This study suggested that BGISEQ-500 is a qualified sequencing platform for WES.

## Background

The launch of the Roche 454 sequencer [1] opened the era of next-generation sequencing (NGS). Compared with the traditional Sanger sequencing technology[2], NGS has significantly larger throughput and lower per-base cost. Taking these advantages, researchers can analyze the information of all the genes in one project, rather than doing gene-by-gene studies. Researchers can obtain the information on the whole genome, protein-coding exons, or other specified regions by performing whole genome sequencing (WGS), whole exome sequencing (WES)[3] or target region sequencing (TRS), respectively. As an easy to interpret, known functional impacts, and relatively low-cost technology comparing with WGS, WES is widely used in human genetics research nowadays.

In the short history of NGS era, five major sequencing platforms have emerged: Roche 454, Illumina Hiseq series (GA, Hiseq, Miseq, X)[4], SOLiD[5], Complete Genomics[6], and Ion Torrent[7]. These platforms use different mechanics and have their specific advantages and disadvantages [8]. After years of technology evolution and competition, Hiseq becomes the most widely used sequencing platform. In 2015, BGI and Complete Genomics jointly announced a new next-generation sequencer, BGISEQ-500. However, its performance on WES has not yet been well evaluated by the scientific community.

We evaluated the performance of BGISEQ-500 on WES by parallel comparison with Hiseq 4000 on the well-characterized human sample NA12878. We compared the concordance of variation detected between the sequencing platforms, and their variation detection accuracy with the reference variation dataset from the genome in a bottle project(GIAB)[9]. We found that BGISEQ-500 has comparable reproducibility and competitive variation detection accuracy to Hiseq 4000.

## Results

### Data Production

The DNA of NA12878 was used in this study. Agilent SureSelect Kit v5 (50.4 Mb) was used for exome capture. The sequencing strategy was pair-end 150bp for Hiseq4000 and pair-end 100bp for BGISEQ-500. The DNA was sequenced to >100x on both Hiseq 4000 and BGISEQ-500 platform. (Methods) Each platform sequenced four replicable libraries, resulting in eight datasets in total. For comparison, each dataset was down-sampled to 100x. BGISEQ showed higher exome capture efficiency (72% vs. 58%), therefore it requires less sequencing data to reach the same sequencing depth. BGISEQ also showed slightly lower duplication rate than Hiseq (7.0% vs 7.6%). For each dataset, >99.6% bases of the target region are covered by at least one read, and >96% bases of the target region are covered with >= 20 reads, indicating that the whole target region is comprehensively and uniformly captured on both platforms. (Table 1, Figure 1)

**Table 1.** Data production

|  | Hiseq-1 | Hiseq-2 | Hiseq-3 | Hiseq-4 | BGISEQ-1 | BGISEQ-2 | BGISEQ-3 | BGISEQ-4 |
| --- | --- | --- | --- | --- | --- | --- | --- | --- |
| Read length | PE150 <sup>a</sup> | PE150 | PE150 | PE150 | PE100 | PE100 | PE100 | PE100 |
| Raw data/Gb | 10.04 | 9.83 | 9.78 | 9.00 | 7.81 | 7.90 | 7.65 | 7.56 |
| Mean depth | 99.77 | 99.78 | 100.37 | 99.66 | 102.27 | 101.88 | 101.92 | 101.77 |
| Capture efficiency <sup>b</sup> (%) | 56.02 | 56.84 | 58.79 | 62.43 | 71.54 | 70.22 | 72.65 | 73.20 |
| Duplication rate | 7.12 | 6.59 | 8.79 | 7.92 | 7.10 | 7.00 | 7.18 | 6.75 |
| Coverage (%) | 99.74 | 99.66 | 99.72 | 99.75 | 99.82 | 99.83 | 99.82 | 99.82 |
| 4x coverage (%) | 99.51 | 99.37 | 99.43 | 99.49 | 99.63 | 99.64 | 99.62 | 99.61 |
| 10x coverage (%) | 98.89 | 98.67 | 98.60 | 98.74 | 98.89 | 98.93 | 98.85 | 98.84 |
| 20x coverage (%) | 97.10 | 96.80 | 96.11 | 96.16 | 96.29 | 96.30 | 96.01 | 96.18 |

a. PE, pair-end.
b. Bases aligned on target region / raw data amount

**Figure 1.**
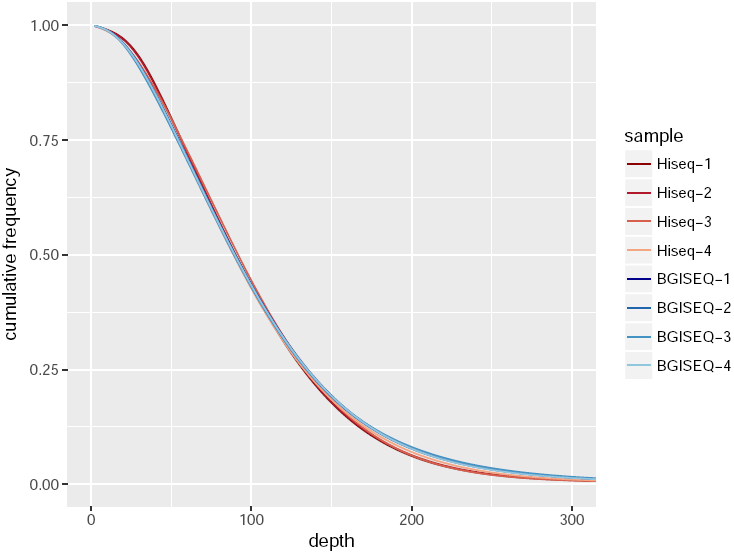
Cumulative depth distribution. The cumulative frequency is the fraction of target regions covered by given depth or higher.

### Variation detection

The variation detection was processed under the guidelines from the Genome Analysis Toolkit (see Methods for details) [10, 11]. Only the target region was used for variation detection. Roughly 41 thousands single nucleotide polymorphisms (SNPs) were detected from each dataset, including 19 thousand inside the protein coding region, and ∼9.4 thousand could lead to protein change (Table 2). The BGISEQ datasets generated slightly fewer SNPs than Hiseq datasets. About 99.7% of detected SNPs could be found in dbSNP142 [12]. The transition/transversion ratio (Ti/Tv) on whole target region and on the exomic region is 2.56 and 3.09, and the corresponding heterozygous to homozygous variation ratio (het/hom) is 1.64 and 1.52, respectively. (Table 2) These metrics from our datasets are comparable to other sources [9, 13].

**Table 2.** Variation detection and annotation

|  |  | Hiseq-1 | Hiseq-2 | Hiseq-3 | Hiseq-4 | BGISEQ-1 | BGISEQ-2 | BGISEQ-3 | BGISEQ-4 |
| --- | --- | --- | --- | --- | --- | --- | --- | --- | --- |
| SNP | total number <sup>a</sup> | 41554 | 41506 | 41627 | 41540 | 41264 | 41294 | 41292 | 41172 |
|  | found in dbSNP(%) | 99.75 | 99.73 | 99.70 | 99.78 | 99.74 | 99.76 | 99.76 | 99.80 |
|  | homozygous | 15741 | 15723 | 15723 | 15758 | 15671 | 15666 | 15692 | 15642 |
|  | heterozygous | 25813 | 25783 | 25904 | 25782 | 25593 | 25628 | 25600 | 25530 |
|  | Ti/Tv | 2.56 | 2.57 | 2.56 | 2.56 | 2.56 | 2.56 | 2.56 | 2.56 |
|  | het/hom <sup>b</sup> | 1.64 | 1.64 | 1.65 | 1.64 | 1.63 | 1.64 | 1.63 | 1.63 |
|  | intronic variations <sup>c</sup> | 17298 | 17351 | 17321 | 17288 | 17217 | 17232 | 17220 | 17229 |
|  | exonic variations | 21540 | 21497 | 21561 | 21533 | 21408 | 21407 | 21433 | 21322 |
|  | coding variations | 19353 | 19332 | 19354 | 19364 | 19269 | 19273 | 19291 | 19210 |
|  | nonsynonmous | 9446 | 9439 | 9437 | 9466 | 9393 | 9391 | 9400 | 9343 |
|  | Ti/Tv on exome | 3.09 | 3.10 | 3.08 | 3.09 | 3.08 | 3.08 | 3.08 | 3.09 |
|  | het/hom on exome | 1.52 | 1.52 | 1.52 | 1.52 | 1.52 | 1.52 | 1.52 | 1.52 |
| indel | total number | 3461 | 3436 | 3470 | 3445 | 3503 | 3559 | 3506 | 3538 |
|  | found in dbSNP(%) | 94.42 | 95.08 | 94.55 | 94.83 | 94.78 | 94.44 | 94.69 | 94.04 |
|  | homozygous | 1491 | 1491 | 1492 | 1502 | 1433 | 1444 | 1432 | 1420 |
|  | heterozygous | 1970 | 1945 | 1978 | 1943 | 2070 | 2115 | 2074 | 2118 |
|  | het/hom | 1.32 | 1.30 | 1.33 | 1.29 | 1.44 | 1.46 | 1.45 | 1.49 |
|  | intronic variations | 2493 | 2493 | 2518 | 2496 | 2558 | 2585 | 2553 | 2581 |
|  | exonic variations | 703 | 689 | 702 | 694 | 705 | 706 | 703 | 702 |
|  | coding variations | 460 | 465 | 459 | 464 | 473 | 477 | 468 | 473 |
|  | het/hom on exome | 1.12 | 1.11 | 1.11 | 1.11 | 1.17 | 1.21 | 1.14 | 1.17 |

a. Only variants on target region were used in these statistics.
b. het/hom, heterozygous to homozygous variation ratio
c. Variations located at splicing sites are considered as nonsynonymous and not count as intronic

Roughly 3.5k insertion/deletions (indels) were detected, approximately 470 out of which lie on coding region. BGISEQ detected slightly more heterozygous indels than Hiseq, resulting in higher het/hom ratio (1.46 vs. 1.31). (Table 2) Around 95% of indels have been previously reported in dbSNP142. The indels from these datasets also showed similar length distribution. (Figure 2)

**Figure 2.**
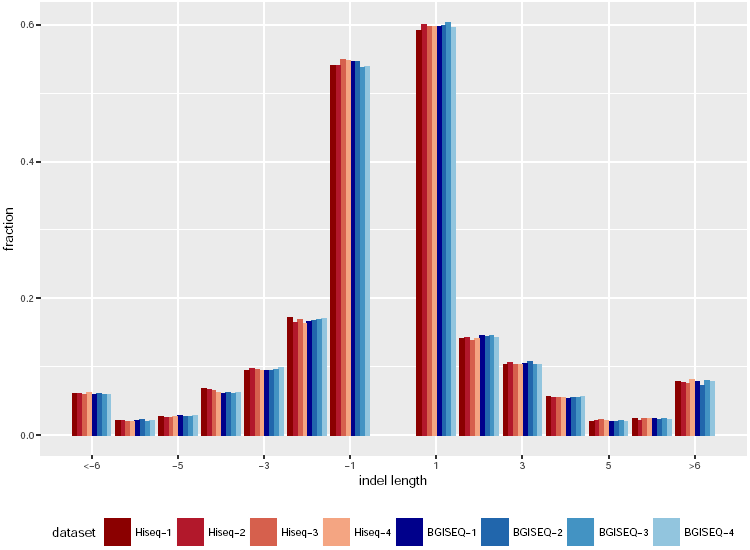
Indel length distribution. Deletions are shown as negative length whereas insertions are shown as positive. The fraction of insertions and deletions sum up to 1 separately. All datasets showed similar length distribution.

### Variation Concordance

It has been noticed that the repetitive sequence in the genome could lead to ambiguity of short fragment alignment, which subsequently leads to false variation detection results. This could be a major cause of the SNP detection errors.[14] Using the genome mappability score[14], ∼2.3% of the target region was identified with alignment uncertainty. These regions were eliminated, and only the mappable regions were used hereafter.

The SNPs and indels from these four datasets were compared against each other separately, and Jaccard similarity was used to measure the concordance between datasets. It is showed that SNP results have 97.6% intra-platform concordance and 96.7% inter-platform concordance, and BGISEQ has slightly higher intra-platform concordance than Hiseq. (Figure 3) The high intra-platform concordance indicated qualified reproducibility of each platform. Moreover, BGISEQ has supreme inter-platform concordance with Hiseq, suggesting that BGISEQ could substitute Hiseq in many application fields where SNPs are the primary focus.

**Figure 3.**
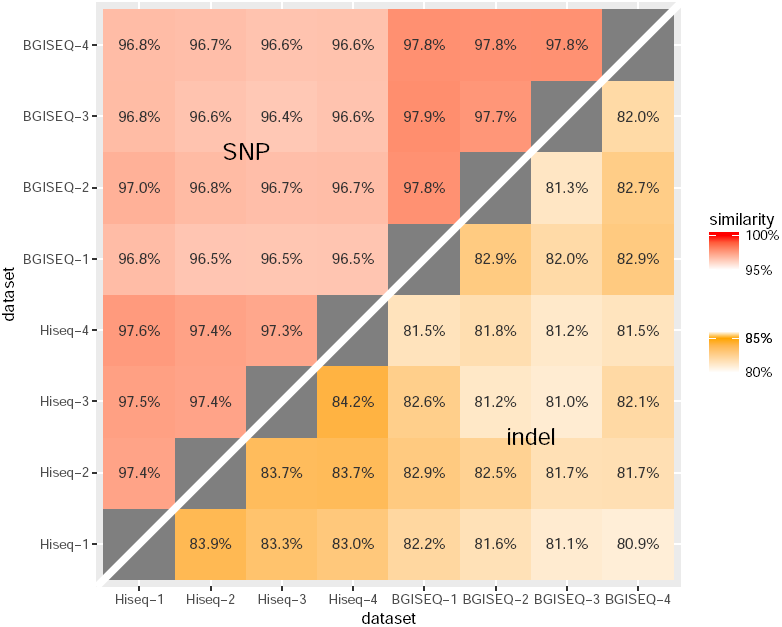
Concordance of variation detection. The Jaccard similarity for variation detection results from datasets was calculated for SNP (top-left triangle) and indel (bottom-right triangle) separately. SNP detection showed excellent intra- and inter-platform concordance, while indel detection showed inferior concordance. The inter-platform concordance is slightly lower than intra-platform concordance.

For indel, the intra-platform concordance is 82.3% for BGISEQ and 83.6% for Hiseq, and the inter-platform concordance is 81.7%. Indels with exact the same position and alter alleles were considered as concordant, regardless of their genotypes. This concordance level is lower than SNP’s, as expected, because it is harder to detect indels from short reads using current methods[15]. It is possible that there could be different concordance metrics for indels, depending on how position and genotype concordance is defined. When only the location is considered, regardless of the genotypes, the intra-platform concordance increased to 87.7% for BGISEQ and 87.4% for Hiseq, and the inter-platform concordance increased to 85.1%. On the other hand, if we restrict concordance sites as exact genotype match, the intra-platform concordance decreased to 79.4% for BGISEQ and 81.3% for Hiseq, and the inter-platform concordance decreased to 76.5%. These results suggest that different datasets and platforms have better agreement on indel location, but diverge on zygosity and genotypes.

### Variation accuracy

As the sequenced sample NA12878 had been well characterized by genome in the bottle project (GIAB) [9], the genotype result from GIAB was used as the reference to benchmark variants accuracy. Only the high confidence region from the GIAB dataset was used for evaluation. Sensitivity and precision were used during the evaluation. Variation detected by both test data and GIAB were considered as true positive if they have exactly the same positions, regardless of whether they have the same genotypes. Overlapping indel positions were not considered as the same one unless they have the same start and end positions.

Regarding SNP detection, 35,210 SNPs were found in the GIAB dataset, and the sensitivity and precision from datasets is 99.0% and 99.4%, respectively. Both platforms have excellent SNP detection accuracy. Furthermore, within 196 false positive and 356 false negative sites per sample, 126 (64%) and 204(57%) are concordant in all 8 samples, respectively, indicating errors in the reference set or systemic exome sequencing bias. Taking this into consideration, the actual accuracy from these platforms should be higher, and the difference between datasets on these loci is minimal.

**Table 3.** Variation accuracy estimation by comparison with GIAB

|  |  | detected SNPs |  |  | GIAB-specific variations | Sensitivity (%) | Precision (%)<sup>a</sup> | F-measure (%)<sup>b</sup> |
| --- | --- | --- | --- | --- | --- | --- | --- | --- |
|  |  | total | in GIAB | not in GIAB |  |  |  |  |
| SNP | Hiseq-1 | 35051 | 34851 | 200 | 359 | 98.98 | 99.43 | 99.20 |
|  | Hiseq-2 | 35026 | 34793 | 233 | 417 | 98.82 | 99.33 | 99.07 |
|  | Hiseq-3 | 35030 | 34821 | 209 | 389 | 98.90 | 99.40 | 99.15 |
|  | Hiseq-4 | 35037 | 34842 | 195 | 368 | 98.95 | 99.44 | 99.20 |
|  | BGISEQ-1 | 35073 | 34883 | 190 | 327 | 99.07 | 99.46 | 99.26 |
|  | BGISEQ-2 | 35069 | 34876 | 193 | 334 | 99.05 | 99.45 | 99.25 |
|  | BGISEQ-3 | 35071 | 34886 | 185 | 324 | 99.08 | 99.47 | 99.28 |
|  | BGISEQ-4 | 35048 | 34881 | 167 | 329 | 99.07 | 99.52 | 99.29 |
|  | Hiseq-1 PE100 | 35110 | 34905 | 205 | 305 | 99.13 | 99.42 | 99.27 |
|  | Hiseq-2 PE100 | 35056 | 34855 | 201 | 355 | 98.99 | 99.43 | 99.21 |
|  | Hiseq-3 PE100 | 35174 | 34880 | 294 | 330 | 99.06 | 99.16 | 99.11 |
|  | Hiseq-4 PE100 | 35143 | 34897 | 246 | 313 | 99.11 | 99.30 | 99.21 |
| indel | Hiseq-1 | 2501 | 2453 | 48 | 197 | 92.57 | 98.08 | 95.24 |
|  | Hiseq-2 | 2493 | 2454 | 39 | 196 | 92.60 | 98.44 | 95.43 |
|  | Hiseq-3 | 2507 | 2457 | 50 | 193 | 92.72 | 98.01 | 95.29 |
|  | Hiseq-4 | 2508 | 2453 | 55 | 197 | 92.57 | 97.81 | 95.11 |
|  | BGISEQ-1 | 2542 | 2480 | 62 | 170 | 93.58 | 97.56 | 95.53 |
|  | BGISEQ-2 | 2571 | 2498 | 73 | 152 | 94.26 | 97.16 | 95.69 |
|  | BGISEQ-3 | 2553 | 2478 | 75 | 172 | 93.51 | 97.06 | 95.25 |
|  | BGISEQ-4 | 2564 | 2488 | 76 | 162 | 93.89 | 97.04 | 95.44 |
|  | Hiseq-1 PE100 | 2538 | 2498 | 40 | 152 | 94.26 | 98.42 | 96.30 |
|  | Hiseq-2 PE100 | 2512 | 2471 | 41 | 179 | 93.25 | 98.37 | 95.74 |
|  | Hiseq-3 PE100 | 2541 | 2474 | 67 | 176 | 93.36 | 97.36 | 95.32 |
|  | Hiseq-4 PE100 | 2523 | 2484 | 39 | 166 | 93.74 | 98.45 | 96.04 |

- a. Precision = true positive / (true positive + false positive). Precision instead of specificity was used because true negative dominate the region thus specificity is very close to 1. - b. F-measure is the harmonic average of the sensitivity and precision. It combines sensitivity and precision in a single measurement.

For indel detection, 2,650 indels are found in the GIAB dataset, and the sensitivity is 93.8% and 92.6%, and the precision 97.2% and 98.0% for BGISEQ and Hiseq, respectively. BGISEQ showed higher sensitivity but lower precision than Hiseq. Thirteen false positive and 42 false negative sites are concordant across samples, contributing to 22.9% and 23.6% total false positives and false negatives on average. To achieve better indel detection performance is still challenging for exome sequencing and GATK pipeline.

### Impact of sequence depth and read length

To analyze the sequence depth effects to variation detection, the raw data were down-sampled to various sequence depths (20x, 30x, 50x, 70x, 100x, 150x). The SNP detection sensitivity increases with increased sequence depth, and it plateaus after the sequence depth exceeds 100x. (Figure 4) The increase in sensitivity may be due to better coverage of the target with increased sequence depth. On the other hand, the SNP detection precision stays constant while sequence depth increased, showing that the model has reached its limit when the depth is greater than 20x. For indel detection, the sensitivity increases while sequence depth increasing, as expected. But it does not reach a plateau even when the sequence depth is as high as 150x, showing that additional data is required for a better indel detection.

**Figure 4.**
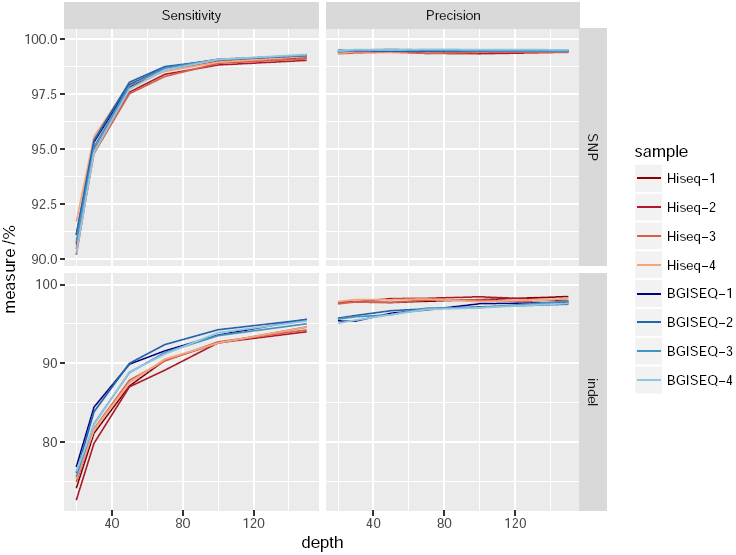
Variation detection accuracy versus sequence depth. Raw data were down-sampled to 20x, 30x, 50x, 70x, 100x, and 150x to generate this plot. Variations on the high confidence region from the genome in the bottle project were used as the reference.

By truncating read length to 100bp, four additional Hiseq pair-end 100bp (PE100) datasets were generated from Hiseq datasets respectively, with the same sequence depth used in the above evaluation. Compared to Hiseq PE150 datasets, PE100 datasets showed similar precision and sensitivity on SNP detection. On indel detection, PE150 and PE100 have similar precision while PE100 showed slightly higher sensitivity (Table 3). The result suggests that the read length has no significant impact on variation detection accuracy at this sequencing depth level.

## Discussion

The reproducibility of a method and its consistency with other available methods are essential for its application in academic and clinical scenarios. The aim of this study was to evaluate these characteristics of BGISEQ-500 in WES studies. Considering that WES is widely used in human genetics research and approaching clinical use, the validation of this newly established platform is crucial.

By comparison with Hiseq4000 data, this study showed BGISEQ could achieve comparable coverage to Hiseq for exome capture and sequencing procedure. For variation detection, both platforms have high and comparable reproducibility, although the reproducibility of indels is lower than of SNPs. Furthermore, the inter-platform concordance is commensurate with intra-platform concordance on SNP detection. This indicates that BGISEQ WES is capable of the applications which Hiseq WES SNP data has been tested and verified for.

For data usage, BGISEQ showed higher exome capture efficiency, and requires about 25% less data than Hiseq to reach the same sequencing depth. If the sequencing cost per gigabase is comparable between the two platforms, BGISEQ will have a lower cost per sample to obtain the same amount of effective data. Furthermore, smaller dataset also means less computational resources and runtime for bioinformatics analysis.

Unlike former Complete Genomics sequencers, the raw data generated from BGISEQ-500 is in fastq format, the *de facto* standard format for NGS data. As a result, the data is acceptable to most of the commonly used analysis software. This allows scientists to manipulate the data by themselves, to adapt the cutting edge analysis methods, and to compare results with other data easily. As an illustration, the commonly used bwa-GATK pipeline was applied to both Hiseq and BGISEQ data seamlessly in this study, keeping the results clear of biases possibly introduced by different analysis software.

It is important to note that BGISEQ is still in its rapidly evolving phase and the amount of data generated from BGISEQ is limited. Therefore, the results showed in this study could be limited by its sample size and the current state of the instruments. The full evaluation and validation of this platform requires more data from research and clinical scenarios.

## Conclusion

In this study, we evaluated the performance of BGISEQ-500 on WES and compared it with Hiseq4000 WES and GIAB high-confidence variation dataset. BGISEQ showed high reproducibility and concordance in intra‐ and inter‐ sequencing platforms. Both BGISEQ and Hiseq platforms demonstrated adequate variation detection accuracy on the benchmark region. These results suggest that BGISEQ-500 is a qualified sequencing platform for WES.

## Methods

### Data Production

The DNA of NA12878 was acquired from Coriell Institute (Catalog ID NA12878). Agilent SureSelect Kit v5 was used for exome capture. The library construction and sequencing procedure on BGISEQ-500 were as described in BGISEQ sequencing section, with a ∼170bp insert size and pair-end 100bp sequencing strategy. The procedure on Hiseq4000 followed the manufacturer instructions with a 250∼300bp insert size and pair-end 150bp sequencing strategy. The DNA was sequenced to >100x on both platforms. Each platform sequenced four replicable libraries. Each dataset was randomly down-sampled to 100x for comparison.

### BGISEQ sequencing

#### DNA preparation

1μg DNA (Qubit quantified) was sheared by Covaris and double selected with Ampure XP beads to acquire fragments around 170bp. End repairing, A-tailing and Ad153 index adapter ligation of 50ng size-selected DNA (0.6*0.8 Ampure XP beads double-selection) were performed in a single tube for a total time of 1.75 h, followed by a purification with 50μL of Ampure XP beads and 20μL of TE buffer. After PRE-PCR following the 95°C 3min, (98°C 20s, 60°C 15s, 72°C 30s) 8 cycles, 72°C 10min,4°C hold thermal cycles using KAPA HiFi Hot Start Ready Mix, the product was purified with 1X Ampure XP beads and quantified with Qubit BR ds DNA kit.

#### Exome capture

The hybridization of BGISEQ-500 library was performed according to the Agilent SureSelect protocol with the following optimized parameters: 1000ng purified DNA was used for hybridization, the index block and PCR block of Agilent were replaced by a corresponding Ad153_index block (one for all indexes) and Ad153_PCR block. KAPA HiFi Hot Start Ready Mix was used for Post-PCR following the 95°C 3min, (98°C 20s, 60°C 15s, 72°C 30s) 13 cycles, 72°C 10min, 4°C hold thermal cycle. Post-PCR products were purified with 1X Ampure XP beads and quantified with Qubit BR ds DNA kit. The fragment size distribution was analyzed using the Agilent 2100 Bioanalyzer and DNA 1000 kit.

#### Single strand DNA (ssDNA) circle construction

300ng Post-PCR products were denatured at 95°C for 3min (with heated lid at 105°C) and transferred to 4°C quickly to make a single strand DNA circle (ssDNA circle). After heat denaturation, the splint oligo binds to the adapters on both ends of a single Post-PCR product, guiding both ends of the single strand to adjacent positions. The following ligation reaction using T4 DNA ligase at 37°C for 30min helps connect the adjacent bases on different ends of the single strand with a phosphodiester bond to complete the circularization. An enzyme digestion using Exo I and Exo III at 37°C, 30min was implemented to eliminate uncirculated DNA. The libraries were purified with 168μL of Ampure XP beads and quantified with Qubit BR ssDNA kit. The resulting ssDNA circle is the final library.

#### Make DNA nanoballs (DNBs)

DNBs were generated from the ssDNA circle using rolling circle amplification (RCA) to enhance the fluorescent signals in the sequencing process[6]. Primer mix bind to 6ng ssDNA circles at 95°C for 1 min, 65°C for 1 min, and 40°C for 1 min. 40μL Phi29 DNA polymerase and 4μL SSB was added for RCA reaction at 30° C for 30 min, then quickly transferred to 4°C. The reaction was ended completely by 20μl stop buffer, generating even-sized DNBs which will have similar fluorescent intensity in the sequencing process to ensure the signal chastity. Compared to PCR amplification, RCA has no PCR error accumulation and no PCR bias because the original ssDNA circle is the only template during the entire amplification process [6]. Unlike emulsion or bridge PCR, the rolling-circle amplification does not require precise titration of template concentrations *in situ* and circumvents stochastic inefficiencies.

#### Loading and sequencing

The DNBs were combined with 1/4 volume of DNB loading buffer and an appropriate amount of PBS buffer to a total volume of 140μL, and placed on the loader machine. The DNBs were loaded onto the flow cell in which DNB binding sites are patterned nano-arrays. Sequencing data were generated with pair-end 100bp sequencing strategy on the BGISEQ-500 platform.

### Variation Detection

The variation detection proceeded under the guidelines from Genome Analysis Toolkit (GATK) [10, 11]. Reads were aligned to human reference genome hg19 using bwa-mem (version 0.7.15)[16] with default parameters. The bam files were sorted, merged and library duplications were identified using Picard (https://github.com/broadinstitute/picard, version 2.5.0). After that, GATK (version 3.3) was applied to refine reads around indels, and recalibrate base quality. Variation calling on the capture region was carried out by GATK HaplotypeCaller with ‘‐‐emitRefConfidence GVCF ‐‐variant_index_type LINEAR ‐‐variant_index_parameter 128000’. Additionally, because the BGISEQ library construction protocol introduced more PCR cycles, we used ‘-pcrModel AGGRESSIVE’ for BGISEQ datasets. The gvcfs were then genotyped by GenotypeGVCFs with ‘-stand_call_conf 30 ‐stand_emit_conf 10 ‐allSites’. Raw SNP and indels were extracted by SelectVariants separately. To obtain high quality variants, hard filter was applied using GATK VariantFiltration but with separate criteria for each platform. We found that the GATK recommendation criteria worked well for illumina data (‘QD < 2.0 || FS > 60.0 || MQ <40.0 || MQRankSum < ‐12.5 || ReadPosRankSum < ‐8.0’ for SNPs; and ‘QD < 2.0 || FS > 200.0 || ReadPosRankSum < ‐20.0’ for indels), but it looks poor for the BGISEQ data. This is reasonable because these recommendation were specific based on illumina data, and it is reasonable to assume that BGISEQ data have different characteristics. A different criteria was used for BGISEQ data (‘QD < 2.0 || FS > 60.0 || ReadPosRankSum < ‐8.0’ for SNPs; and ‘QD < 4.0 || FS > 200.0 || ReadPosRankSum < ‐8.0’ for indels). The final call-sets were then annotated by SnpEff (version: 4.0) [17].

### Variation concordance analysis

Regions with genome mappability scores (GMS) [14] less than 1 were excluded from the evaluation. Jaccard similarity (number of sites where both datasets detected as SNP divided by the number of sites where at least one dataset is detected as SNP) was used to measure the concordance between datasets.

### Variation accuracy estimation

The genotype result from GIAB was used as the reference to benchmark variation accuracy. Only the GIAB high confidence region with GMS equal to 1 was used. During the evaluation, precision (true positive / (true positive + false positive)) instead of specificity (true negative / (true negative + false positive)) was used because true negative dominated the dataset. Variation loci detected in both test data and GIAB were considered as true positive.

## Data availability

The datasets generated during the current study are available on CNGB Nucleotide Sequence Archive (CNSA) with project accession CNP0000165.

## Abbreviations

DNB: DNA nanoballs

GATK: Genome Analysis Toolkit

GIAB: genome in a bottle project

GMS: genome mappability score

het/hom: heterozygous to homozygous variation ratio

indel: insertion and deletion

NGS: next-generation sequencing

RCA: rolling circle amplification

SNP: single nucleotide polymorphism

ssDNA: single strand DNA

Ti/Tv: transition/transversion ratio

WES: whole exome sequencing

## Acknowledgments

We thank Ao Chen, Meihua Gong and Kexin Ma from BGI-Shenzhen for their help during the data production on BGISEQ-500, and Scott Gablenz from Complete Genomics for assistance with grammar correction.

## Author contributions

QG, LY, YX, and ZL designed the study. WZ, CX, Jingjing W, HZ, and YT performed the experiment. ZL, YX, and XW analyzed the data. YX, YT, and ZL drafted the paper. YX prepared the figures and tables. CT, LY, YC, XW, Jian W, and HY revised the paper. All the authors approved the paper.

